# Obesity promotes urinary tract infection by disrupting urothelial immune defenses

**DOI:** 10.1101/2025.04.04.647270

**Authors:** Laura Schwartz, Kristin Salamon, Aaron Simoni, Israel Cotzomi-Ortega, Yuriko Sanchez-Zamora, Sarah Linn-Peirano, Preeti John, Juan de Dios Ruiz-Rosado, Ashley R. Jackson, Xin Wang, John David Spencer

## Abstract

Obesity is a significant public health concern that is associated with numerous health risks. Infections are a major complication of obesity, but the mechanisms responsible for increased infection risk are poorly defined. Here, we use a diet induced obesity mouse model and investigate how obesity impacts urinary tract infection (UTI) susceptibility and bladder immune defenses. Our results show that high-fat diet fed female and male mice exhibit increased susceptibility to uropathogenic *E. coli* (UPEC) following experimental UTI. Transcriptomic analysis of bladder urothelial cells shows that obesity alters gene expression in a sex-specific manner, with distinct differentially expressed genes in male and female mice, but shared activation of focal adhesion and extracellular matrix signaling. Western blot and immunostaining confirm activation of focal adhesion kinase, a central component of the focal adhesion pathway, in the bladders of obese female and male mice. Mechanistically, experiments using primary human urothelial cells demonstrate that focal adhesion kinase overexpression promotes UPEC invasion. These findings demonstrate that obesity enhances UTI susceptibility by activating focal adhesion kinase and promoting bacterial invasion of the urothelium. Together, they explain how obesity promotes UTI vulnerability and identify modifiable targets for managing obesity-associated UTI.

**Significance Statement:** Obesity is associated with an increased risk of urinary tract infections (UTIs), but the underlying mechanisms promoting infection susceptibility remain poorly understood. Here, we show that diet-induced obesity drives sex-specific changes in bladder urothelial gene expression, including distinct immune responses in male and female mice. Despite these differences, both sexes exhibit activation of focal adhesion kinase (FAK). FAK overexpression promotes bacterial invasion into human bladder cells. These findings provide a mechanistic explanation for obesity-associated UTI susceptibility and suggest that targeting FAK signaling could offer a therapeutic strategy to prevent UTIs, with implications for personalized interventions in obesity.

## INTRODUCTION

Obesity is a major public health challenge affecting over a billion people globally (1). Obesity can lead to chronic health issues, including type 2 diabetes, cardiovascular disease, nonalcoholic fatty liver disease, neurodegenerative disorders, and increased cancer risk. Beyond its direct health effects, obesity places a significant burden on healthcare systems, reduces overall productivity, quality of life, and life expectancy (2, 3).

Infections are a significant yet overlooked consequence of obesity. Obesity increases susceptibility to nosocomial, surgical site, and community-acquired infections (4, 5). Among the common community-acquired infections, obesity has been identified as an independent risk factor for urinary tract infections (UTI) in both males and females across the lifespan (4–11). Obese individuals with UTI are 2.5 times more likely to require hospitalization compared to their non-obese counterparts (12, 13). Among hospitalized patients, those with obesity are more likely to develop UTI, which can prolong hospital stays (4, 14–16). Obese individuals are nearly five times more likely to develop pyelonephritis, with young girls and women at highest risk (6, 17). Additionally, in kidney transplant recipients, for whom UTI is a significant risk factor for allograft loss, obesity is a significant risk factor for UTI, with obese patients having 4.7 times higher UTI risk than non-obese patients (18). Beyond acute and severe UTIs, obesity is associated with recurrent infections, underscoring its broader impact on urinary tract health (7). Together, these clinical outcomes reveal a gap in our understanding of how obesity influences UTI pathogenesis and highlight the need to define mechanisms driving increased infection risk.

Uropathogenic *Escherichia coli* (UPEC) is the leading cause of UTIs (19). The bladder employs a multifaceted defense system to prevent UPEC infection. In the bladder, the urothelium serves as a critical physical barrier that resists UPEC invasion, reinforced by tightly joined umbrella cells, a uroplakin-and glycosaminoglycan-coated surface, and tight junctions (20). Additionally, the urothelium secretes antimicrobial peptides that directly kill bacteria. Upon pathogen detection, the urothelium activates pattern recognition receptors, triggering downstream transcription factors that promote the expression of antimicrobial peptides as well as cytokines or chemokines. These immune mediators recruit myeloid cells to clear UPEC (20–23).

Despite this integrated urothelial defense network to prevent UTI, UPEC have evolved mechanisms to invade and persist within the bladder. Using adhesive organelles called type 1 pili, UPEC bind to host receptors and are internalized by urothelial cells. Once inside, UPEC can evade host immune responses and resist antibiotics. These intracellular niches may contribute to UTI severity, particularly when urothelial defenses are impaired (23, 24). However, how obesity influences bladder immunity and promotes UTI pathogenesis remains poorly understood.

This study investigates how obesity affects urothelial antibacterial defenses and impacts UTI risk. We utilize a diet-induced obesity mouse model that mirrors human obesity in several ways – including weight gain, insulin resistance, and altered lipid metabolism (25, 26). Mice are fed a high-fat diet (HFD), consisting of 45-60% of calories from fat, to induce obesity. The metabolic abnormalities observed in HFD-fed mice closely parallel human obesity progression, with early weight gain by 2-4 weeks and marked metabolic dysfunction by 12-16 weeks (25, 26). Because male mice develop more severe obesity and metabolic impairment than females, we also assess sex-specific differences in obesity-associated immune and urothelial responses (25–28). To enhance translational relevance, our preclinical findings are validated in primary human urothelial cultures. Using these complementary models, we show that obesity increases UPEC susceptibility via activation of focal adhesion kinase (FAK) signaling in male and female mice. FAK overexpression promotes UPEC invasion, revealing a mechanistic pathway through which obesity augments UTI vulnerability.

## RESULTS

### Obesity increases UTI susceptibility in female mice

To determine whether obesity impacts UPEC vulnerability, female mice were fed either standard chow (SC) or a HFD for 4, 8, or 12 weeks. These timepoints allow us to study the early (4 weeks), progressive (8 weeks), and sustained effects of obesity (12 weeks). After 4 weeks, HFD-fed mice exhibited an 18% increase in body weight compared to SC mice, along with mild elevations in blood glucose concentrations and impaired glucose disposal with glucose tolerance testing. These metabolic disturbances became more pronounced at 8 and 12 weeks. By 12 weeks, HFD-fed mice gained 44% more body weight compared to SC-fed mice and developed more pronounced insulin resistance, as assessed by glucose tolerance testing (**Figure 1A-C**). Female mice did not meet criteria for diabetes, defined by non-fasting blood glucose concentrations ≥ 250 mg/dL or the presence of glucosuria (29). To assess systemic inflammation, we measured serum concentrations of proinflammatory cytokines, but detected no significant differences between SC- and HFD-fed mice (**Supplemental Table 1**).

**Figure 1:**
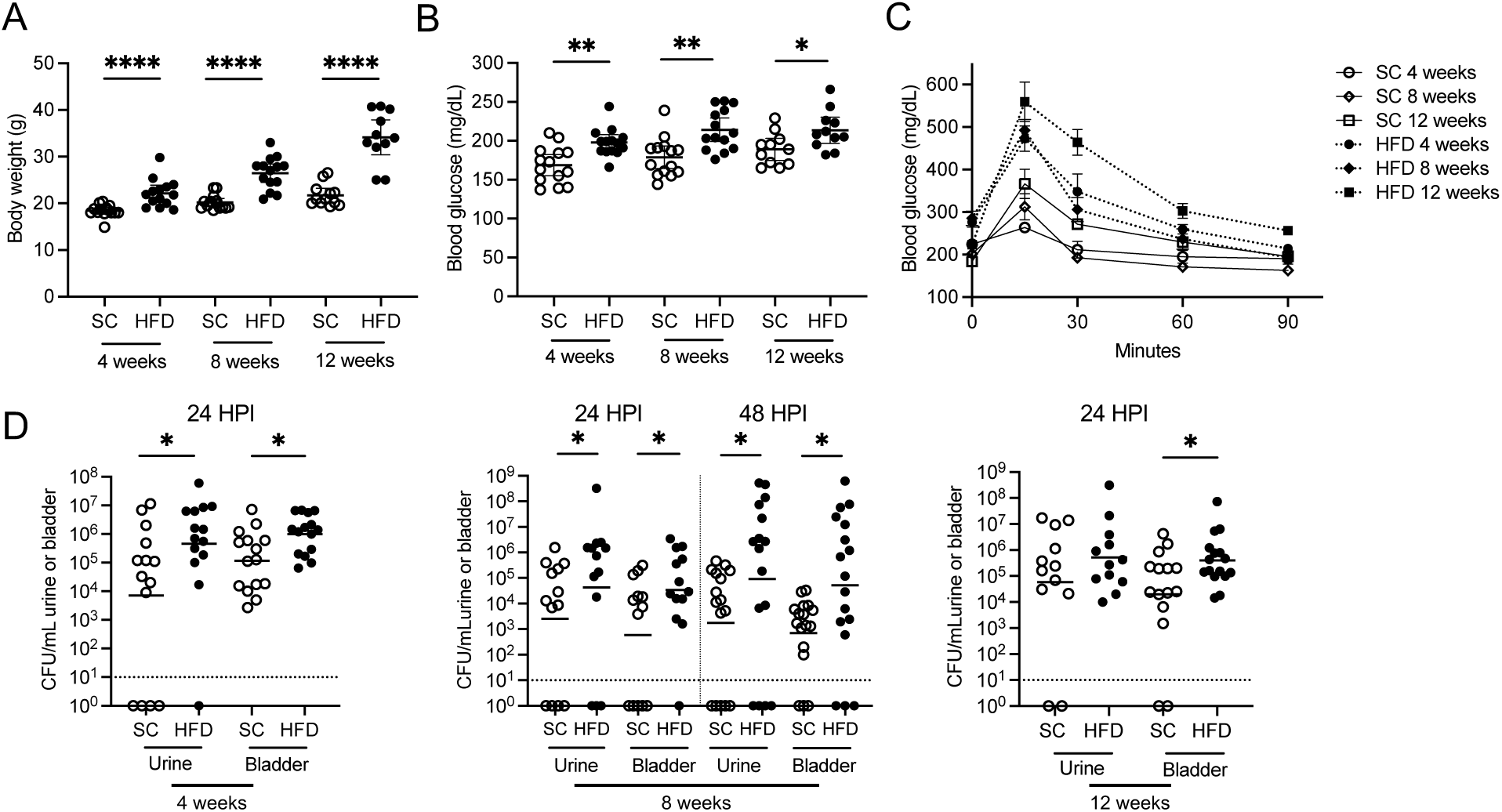
Obesity increases susceptibility to uropathogenic *E. coli* in female mice. (**A-C**) Body weight (A), non-fasting blood glucose (B), and blood glucose concentrations following intraperitoneal glucose tolerance testing (C) measured in female C57/BL6 mice maintained on standard chow (SC) or a high-fat diet (HFD) for 4, 8, or 12 weeks. Graphs show the mean values and SEM. (**D**) Urine and bladder UPEC (UTI89) burden enumerated after transurethral UTI at the indicated time points. HPI notes hours post infection. The horizontal line indicates the geometric mean of each group. The dotted line indicates the limits of detection. (A-D) Each data point represents a measurement from an individual animal. Asterisks indicate significant *P-*values for the indicated pairwise comparison (Mann-Whitney *U* test). \**P* <0.05, \*\**P* <0.01, and \*\*\*\**P* <0.0001.

Given these metabolic impairments, we next assessed whether obesity influences UTI susceptibility. Twenty-four hours after transurethral UPEC infection, HFD-fed mice exhibited greater urine and bladder UPEC titers compared to SC mice (**Figure 1D**). Notably, UPEC burden differences increased with the duration of HFD exposure. Compared to SC fed mice, mice receiving a HFD for 4 weeks had a 2.4-fold increase in mean bladder UPEC CFUs. After 8 and 12 weeks, mean bladder CFUs increased 6.4-fold 24 hours after infection. To determine whether UPEC infection persisted, we enumerated urine and bladder UPEC colonies 48 hours post-infection in mice maintained on HFD or SC for 8 weeks. HFD-fed mice had sustained elevations in UPEC titers compared to SC-fed controls (**Figure 1D**).

### Obesity also increases UPEC vulnerability in male mice

Because sex affects diet-induced obesity outcomes, we next evaluated UTI susceptibility in male mice fed either a HFD or SC. By 8 weeks on a HFD, male mice gained 24% more weight compared to SC-fed mice, and by 12 weeks, HFD-fed mice gained 48% more weight compared to SC controls (**Figure 2A**). Compared to female mice, metabolic impairments were more pronounced in males, with significant elevations in blood glucose concentrations and impaired glucose disposal on glucose tolerance testing, particularly after 12 weeks on a HFD (**Figure 2B/C**). Like female mice, HFD-fed male mice did not develop glucosuria and had comparable serum cytokine concentrations (**Supplemental Table 1**). Following direct bladder UPEC infection, HFD-fed male mice had greater urine and bladder UPEC titers at both 24- and 48-hours post-infection (**Figure 2D**). By 8 and 12 weeks, mean bladder UPEC CFUs were 2.5-fold higher in HFD-fed male mice compared to SC-fed mice 24 hours after infection, while mean urine burden increased 2-fold and 970-fold, respectively. These findings demonstrate that obesity enhances UTI susceptibility in both male and female mice.

**Figure 2:**
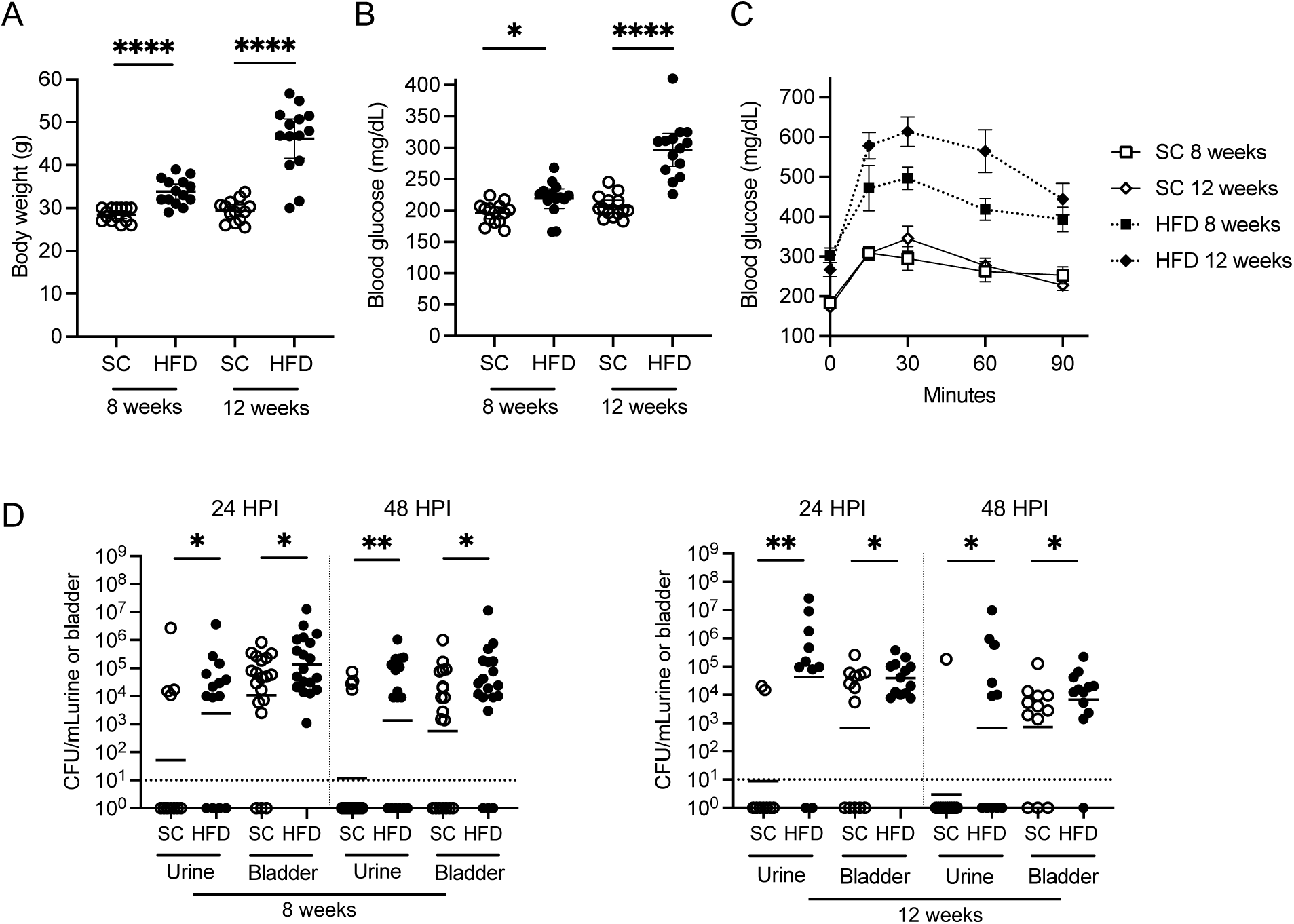
Obesity increases susceptibility to uropathogenic *E. coli* in male mice. (**A-C**) Body weight (A), non-fasting blood glucose (B), and blood glucose concentrations following intraperitoneal glucose tolerance testing (C) measured in male C57/BL6 mice maintained on standard chow (SC) or a high-fat diet (HFD) for 8 or 12 weeks. Graphs show the mean values and SEM. (**D**) Urine and bladder UPEC (UTI89) burden enumerated after direct bladder inoculation at the indicated time points. HPI notes hours post infection. The horizontal line indicates the geometric mean of each group. The dotted line indicates the limits of detection. (A-D) Each data point represents a measurement from an individual animal. Asterisks indicate significant *P-*values for the indicated pairwise comparison (Mann-Whitney *U* test). \**P* <0.05, \*\**P* <0.01, and \*\*\*\**P* <0.0001.

### Obesity augments UTI susceptibility independent of changes in immune cell recruitment and antibacterial function

Myeloid cells, including macrophages and neutrophils, have essential roles in UPEC clearance by migrating to the site of infection, phagocytosing bacteria, and killing pathogens (22, 23). To determine whether diet-induced obesity impairs their recruitment or function, we used flow cytometry to quantify myeloid populations in UPEC infected bladders from male and female mice fed a HFD or SC for 8 or 12 weeks. Recruitment of neutrophils, monocytes, and both M1-like and M2-like macrophages was comparable between HFD-fed and SC mice. Although differences in infection technique prevent direct sex comparisons, we observed fewer neutrophils, monocytes, and macrophages in male mice (SC and HFD) at 8 weeks compared to female mice. Similar trends were observed for monocytes and M2-like macrophages in the 12-week cohort – consistent with prior data showing greater immune cell infiltration in infected female bladders (**Supplemental Figure 1**) (30).

To assess immune cell function, we evaluated myeloid cell phagocytosis, intracellular UPEC killing, and *ex vivo* bacterial killing by neutrophils. UPEC uptake as well as intracellular and extracellular killing were comparable between diet groups (**Supplemental Figures 2 and 3**). However, compared to females, phagocytic activity was reduced in male neutrophils, monocytes, and macrophages (**Supplemental Figure 2**) – supporting published data that phagocytic capacity may be impacted by sex (30). Together, these findings suggest that the increased bacterial burden observed in HFD-fed mice is not attributable to defects in immune cell infiltration or antibacterial function.

### Obesity alters gene expression in the bladder urothelium

Since obesity did not impair myeloid cell recruitment or bacterial killing, we examined whether it compromises urothelial defenses. We performed limiting cell RNA-seq on non-infected bladder urothelium from female and male mice fed SC or a HFD for 12 weeks (*n*=4/cohort, **Figure 3**, **Supplemental Dataset 1**). To validate these findings, an additional RNA-seq analysis was performed in a smaller number of non-infected female and male mice maintained on SC or a HFD for 8 weeks (*n*=3/cohort, **Supplemental Figure 4**, **Supplemental Dataset 2**). Principal component analysis (PCA) demonstrated that urothelial gene expression profiles were consistent among mice within the same diet group but differed between HFD-fed and SC mice, with the greatest variance captured along principal component 1 (**Figure 3A/D and Supplemental Figure 4**). Heatmaps of the differentially expressed genes (DEGs, false discovery rate < 0.05) demonstrated distinct diet-dependent gene expression patterns in both sexes (**Figure 3B/E and Supplemental Figure 4**). These findings indicate that a HFD induces transcriptional changes in the urothelium, suggesting a urothelial-intrinsic response to metabolic stress.

**Figure 3:**
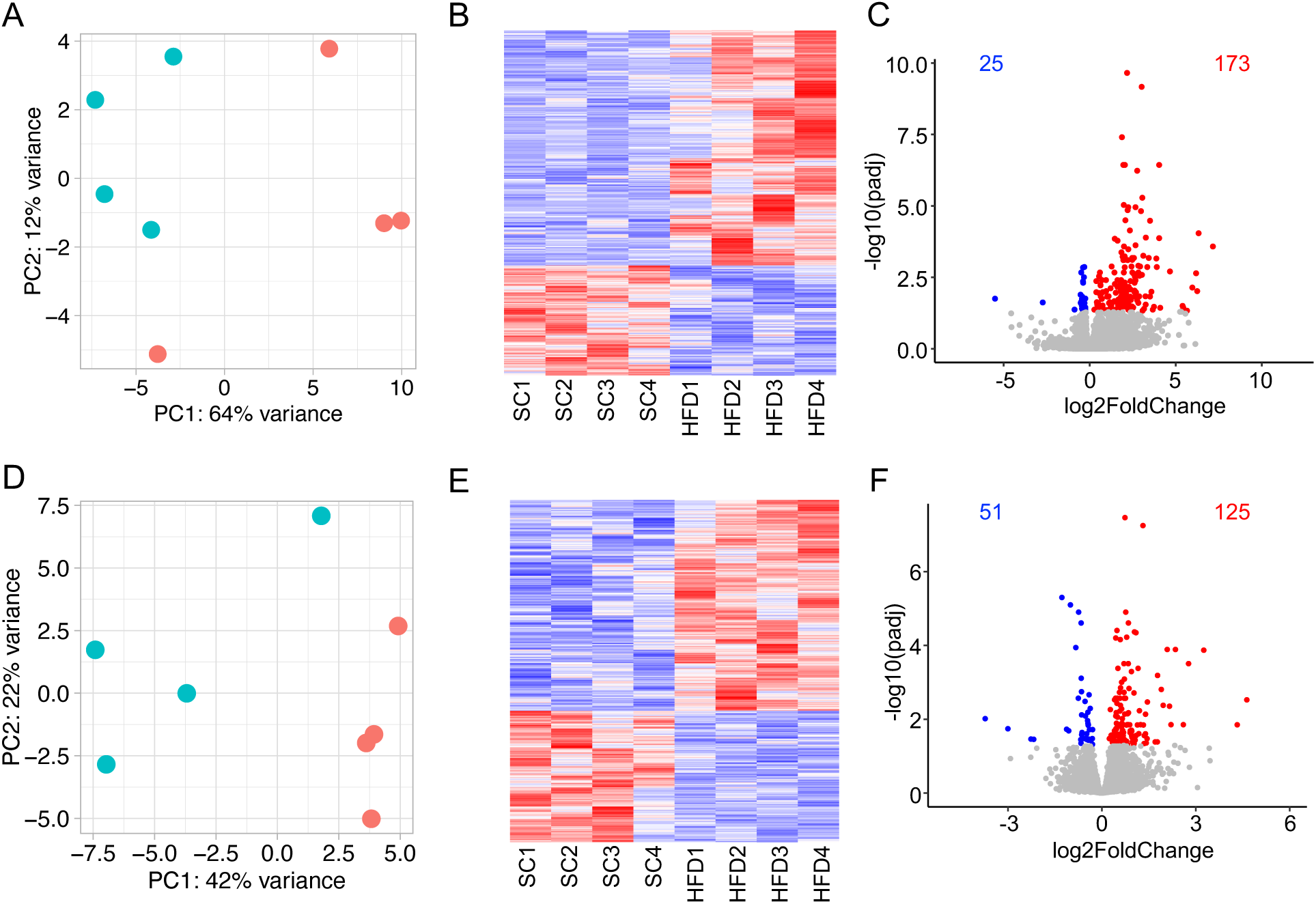
Obesity alters gene expression in the female and male urothelium. (**A/D**) Principal component analysis (PCA) plots from RNA-seq data of female (A) and male (D) urothelium after 12 weeks on standard chow (SC) or high-fat diet (HFD). Each symbol represents a urothelial sample from an individual mouse fed SC (green) or HFD (red). (**B**/**E**) Heatmaps showing differential gene expression between HFD and SC fed female (B) and male (E) urothelium, with each column representing an individual mouse. (**C**/**F**) Volcano plots illustrating differentially expressed genes expression between HFD and SC fed female (C) and male (F) urothelium. Red marks up-regulated genes with log2 fold change > 0 and adjusted *P*-value < 0.05. Blue marks down-regulated genes with log2 fold change < 0 and adjusted *P*-value < 0.05. Adjusted *P*-values were calculated using the Benjamini-Hochberg procedure.

In urothelium from 12-week HFD-fed female mice, 173 genes were up-regulated and 25 genes were down-regulated compared to SC mice (**Figure 3C**). In urothelium from 12-week HFD-fed male mice, 125 genes were up-regulated and 51 genes were down-regulated compared to SC mice (**Figure 3F**). At 8 weeks, urothelium from HFD-fed female mice had 1,812 up-regulated and 1,064 down-regulated genes, whereas HFD-fed male mice had 94 up-regulated and 215 suppressed genes (**Supplemental Figure 4**). The reason for the higher number of DEGs in females after 8 weeks of HFD compared to the 12-week cohort is unclear. While smaller sample sizes often reduce statistical power and result in fewer DEGs, it is possible that this early response may reflect acute transcriptional remodeling or underlying metabolic variability.

### Diet-induced obesity alters urothelial gene expression in a sex-specific manner

Transcriptomic analysis revealed that obesity differentially affects gene expression in female and male urothelium, with minimal overlap in DEGs between sexes. At 12 weeks on a HFD, only two genes were commonly up-regulated in both male and female urothelium, and no genes were commonly down-regulated (**Figure 4A**). A similar pattern was observed at 8 weeks, with just 2 shared up-regulated and 3 shared down-regulated genes (**Supplemental Figure 5**). Despite this minimal gene-level overlap, pathway enrichment analysis showed convergence in the biological processes affected by obesity. Kyoto Encyclopedia of Genes and Genomes (KEGG) pathway enrichment analysis demonstrated that up-regulated genes in HFD mice of both sexes were enriched for pathways related to focal adhesion, extracellular matrix (ECM)-receptor interaction, and cytoskeleton organization (**Figure 4B and Supplemental Figure 5**). Down-regulated genes at 8 and 12 weeks on the diet enriched KEGG pathways related to cancer, however the biological significance of this is unclear due to low enrichment scores and a small number of contributing genes (**Figure 4B and Supplemental Figure 5**). Thus, while obesity drives largely sex-specific changes at the individual gene level, it also induces common transcriptional shifts in pathways involved in cellular architecture and tissue remodeling.

**Figure 4:**
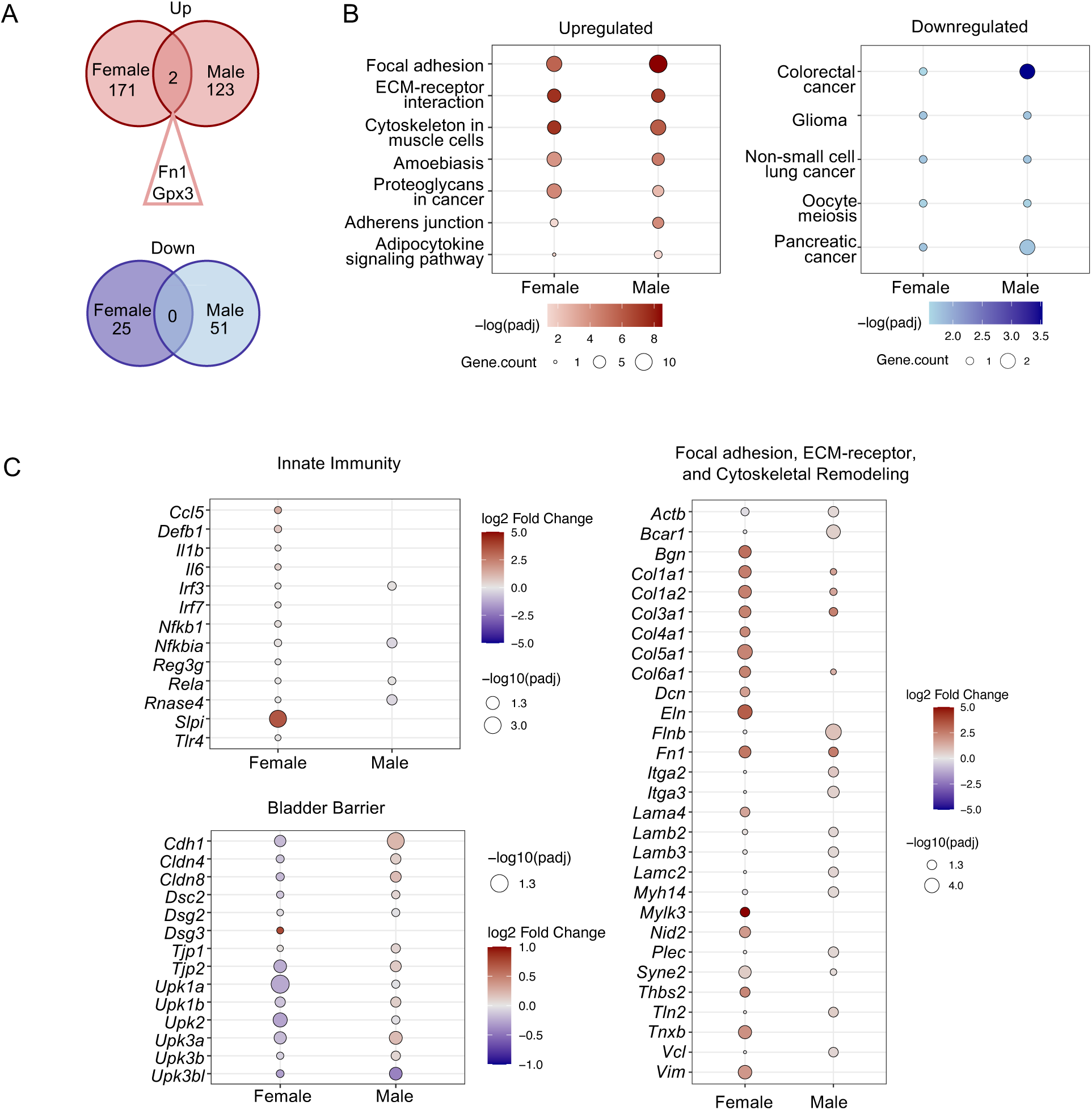
Obesity increases cell-ECM signaling in the urothelium. (**A**) Venn diagrams show minimal overlap between significantly up-regulated (top) and down-regulated (bottom) genes in female and male urothelium after 12 weeks on standard chow (SC) or high-fat diet (HFD). (**B**) KEGG pathway enrichment derived from up-regulated (left) and down-regulated (right) genes in HFD-fed female and male mice. Bubble size reflects the number of enriched genes per pathway and color indicates significance. (**C**) Expression of curated gene sets related to innate immunity, bladder barrier integrity, and focal adhesion/ECM/cytoskeletal remodeling. Bubble color represents log2 fold change (HFD vs. SC) and size indicates significance. (B-C) Significance was assessed via hypergeometric testing on active subnetworks identified from the protein-protein interaction network using PathfindR, with *P*-values adjusted using the Benjamini-Hochberg method.

To investigate how obesity affects established urothelial defenses against UPEC, we profiled the expression of genes involved in innate immunity and epithelial barrier integrity (22, 23). While 12 weeks of HFD had limited impact on innate immune gene expression, genes associated with bladder barrier function and uroplakin plaque formation were disrupted in a sex-specific manner (**Figure 4C**). In females, most bladder barrier genes were suppressed, while in males, many of the same genes were up-regulated. Additionally, genes associated with the KEGG-enriched pathways identified above – including focal adhesion, ECM-receptor interaction, and cytoskeletal organization – were broadly up-regulated in both sexes, with more pronounced changes in HFD-fed female mice (**Figure 4C**). Similar trends were observed at 8 weeks of HFD feeding (**Supplemental Figure 5**). These results suggest that obesity remodels the urothelium, impacting both the apical surface and the cell-ECM interface, potentially compromising the urothelial barrier and increasing susceptibility to UTI.

### Diet-induced obesity activates focal adhesion kinase

Since our transcriptomic analysis revealed enrichment and activation of focal adhesion, ECM-receptor interaction, and cytoskeletal signaling pathways in the urothelium of HFD-fed female and male mice, we next investigated whether obesity alters the urothelial-ECM interface. Focal adhesion kinase (FAK), a non-receptor tyrosine kinase, serves as a central hub for these pathways, integrating signals from the ECM and coordinating downstream intracellular responses, including cytoskeletal remodeling (31, 32). Based on this role, we hypothesized that FAK mediates obesity-driven changes in urothelial structure that promote UPEC infection.

FAK is regulated post-translationally. Upon binding to integrins, it undergoes autophosphorylation and recruits downstream effectors such as PI3K/AKT (31, 32). In urothelium from 12-week HFD-fed female and male mice, Western blot revealed increased FAK and AKT phosphorylation compared to SC controls (**Figure 5A/C).** FAK activation was also observed at 8 weeks of HFD feeding **(Supplemental Figure 6**). Immunostaining confirmed increased FAK expression in the bladder urothelium of mice at both 8 and 12 weeks (**Figure 5B/D and Supplemental Figure 7**). These results corroborate our transcriptomic findings and demonstrate that obesity activates urothelial FAK signaling, which may increase UPEC susceptibility.

**Figure 5:**
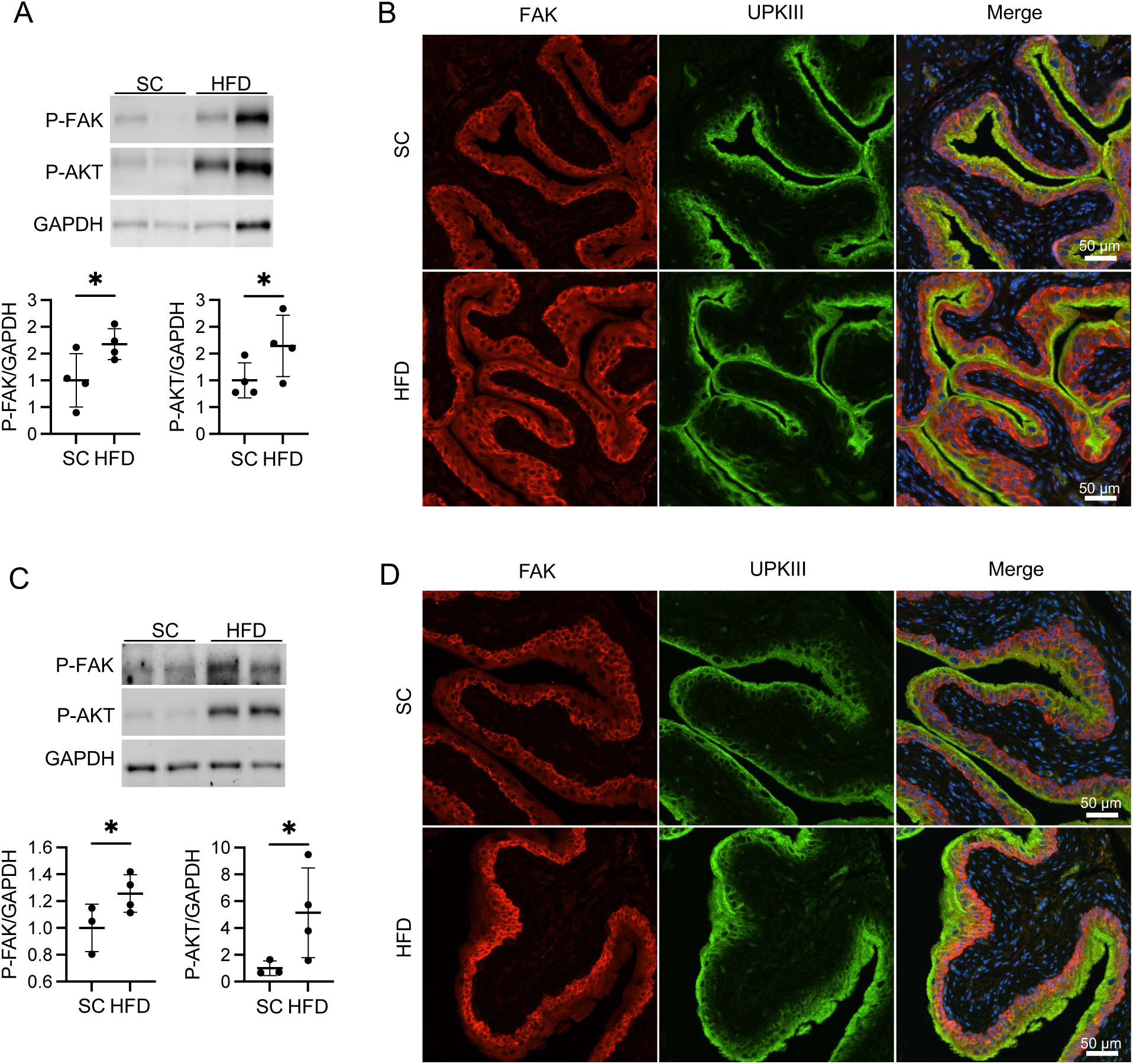
High-fat diet increases focal adhesion kinase activation in the bladder urothelium. (**A**/**C**) Representative Western blots from (A) female and (C) male urothelium after 8 weeks on standard chow (SC) or high-fat diet (HFD) probed for P-FAK (Y397), P-AKT (S473), and GAPDH. Bottom: Densitometry show relative abundance of P-FAK (Y397) and P-AKT (S473), normalized to GAPDH. Each point represents an individual mouse (*n*=4/cohort). Graphs show the mean and standard deviation. Statistical significance was assessed using the Mann-Whitney *U* test. \**P* <0.05. (**B**/**D**) Representative immunofluorescent bladder images from non-infected (B) female and (D) male mice labeled with antibodies to total FAK (red), uroplakin III (green) and nuclei (blue, Hoechst). Magnification = 20X. Staining was completed on 4 bladders per cohort.

### FAK is required and sufficient for UPEC invasion

To further define the role of FAK in UTI, we overexpressed or silenced FAK in primary human urothelial cells and quantified bacterial internalization following UPEC challenge. FAK overexpression significantly increased UPEC invasion, while FAK silencing reduced UPEC entry (**Figure 6A/B**). To test whether this effect is mediated by the downstream effector AKT, we overexpressed AKT and measured UPEC invasion. AKT overexpression had no impact on bacterial uptake, indicating that FAK promotes bacterial entry independent of AKT signaling **(Figure 6C).**

**Figure 6:**
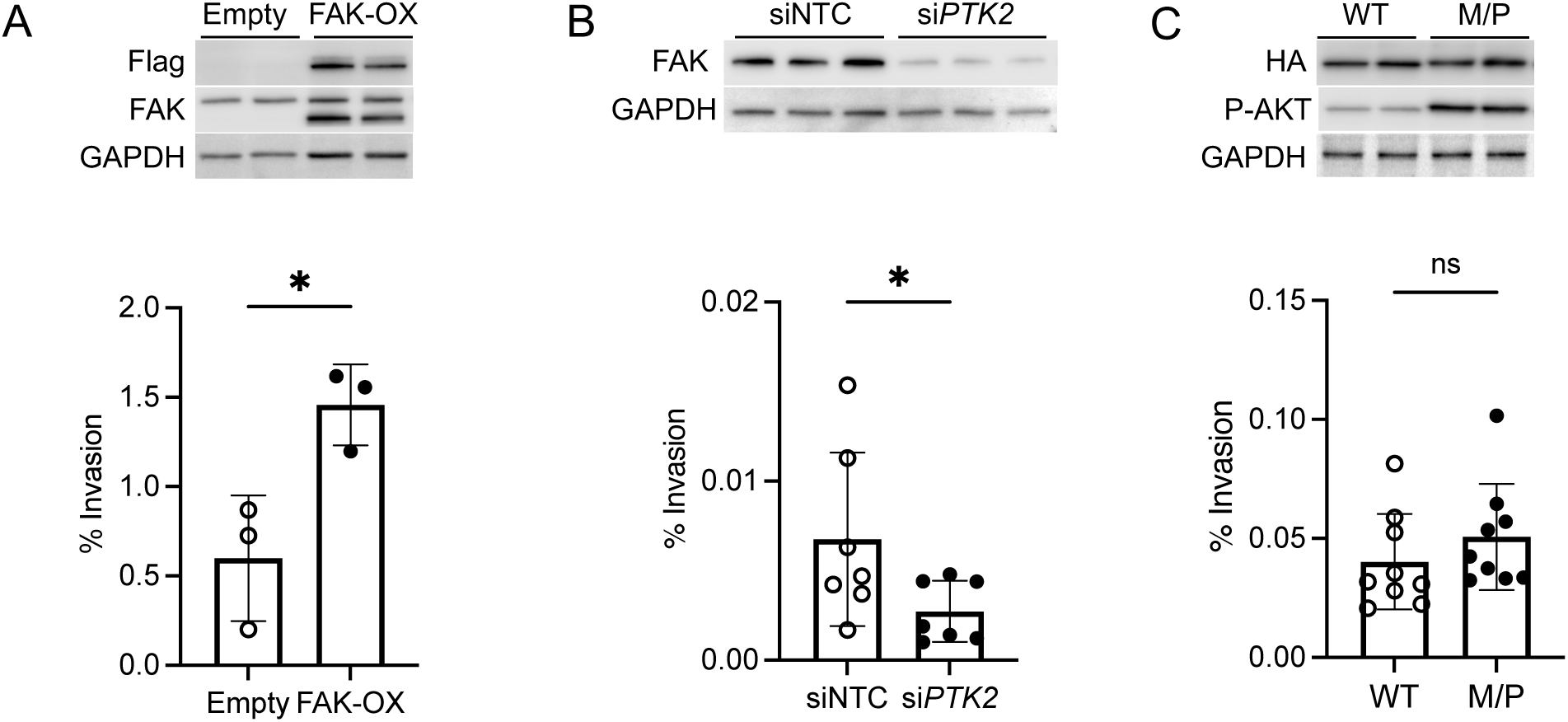
FAK is both required and sufficient for UPEC invasion of human urothelial cells. (**A**) Human bladder urothelial cells were transfected with empty vector (Empty) or Flag-tagged FAK (FAK-OX). Top: Representative Western blot confirmed increased Flag tag expression FAK expression in FAK-OX cells. Bottom: Transfected cells were challenged with UPEC. Shown are the percentage of bacteria invading the cells. (**B**) Human urothelial cells were transfected with non-targeting siRNA control (siNTC) or PTK2-targeting siRNA (siPTK2), which encodes FAK. Top: Representative Western blot confirmed FAK knockdown. Bottom: Transfected cells were challenged with UPEC and the percentage of invading UPEC are shown. (**C**) Human urothelial cells were transfected with HA-tagged wild-type AKT (WT) or membrane-targeted AKT (M/P). Top: Representative Western blot confirmed HA tag expression and increased P-AKT expression. Bottom: Transfected cells were challenged with UPEC and shown and the percentage of invading UPEC are shown. (A-C) Graphs show the mean and SEM. Results are from 3-9 independent experiments performed in triplicate (*n*=3-9). Asterisks indicate significant *P-*values (Student’s *t-*test). \**P* <0.05.

## DISCUSSION

Obesity is an established risk factor for UTI, but the mechanisms underlying this increased susceptibility remain poorly defined (2–4, 27, 33). In this study, we demonstrate that diet-induced obesity increases UTI susceptibility and activates urothelial FAK signaling in both female and male mice. Using human urothelial cells, we show that FAK is both necessary and sufficient to drive UPEC invasion. These findings provide the first mechanistic link between obesity and UTI vulnerability, identifying FAK as a conserved and critical mediator of UPEC susceptibility in the obese bladder.

Our results build on clinical observations connecting obesity to increased UTI incidence and severity (4–11). While chronic inflammation and immune cell dysfunction have been proposed as contributing factors, direct mechanistic evidence is limited (2, 4, 34, 35). In our model, obesity did not impact systemic inflammatory cytokines, impair phagocyte recruitment, or affect UPEC phagocytosis and killing, suggesting that enhanced susceptibility may be driven by other factors. Instead, we observed transcriptional remodeling in the urothelium, implicating uroepithelial vulnerability as a key contributor to UTI pathogenesis.

Obesity has been shown to disrupt epithelial integrity at mucosal surfaces, such as the lung, skin, reproductive tract, and gut (36, 37). In the bladder, we similarly identified sex-specific alterations in genes involved in barrier function and uroplakin plaque formation. In female mice, obesity suppressed several barrier-associated genes and uroplakin transcripts, while in male mice, many of these same genes were up-regulated. One possibility is that female urothelium is more transcriptionally sensitive to dietary fat exposure. Alternatively, male mice, which exhibit more pronounced metabolic dysfunction, may develop maladaptive remodeling programs in response to metabolic stress (2, 38). These findings point to metabolically driven deregulation of bladder urothelial structure and gene programs involved in epithelial maturation and barrier formation, albeit through sex-specific trajectories.

Hormonal signaling may be a key modulator of these effects. Insulin signaling, which is disrupted in obesity, is essential for maintaining epithelial integrity and immune responses (39–43). In prior work, we showed that insulin receptor deletion in the apical urothelium impairs barrier function and increases UTI severity – supporting a model in which metabolic cues influence epithelial remodeling (44). Sex hormones also influence the uroepithelium and UTI risk (45). Estrogen promotes urothelial differentiation and innate defense, while testosterone dampens immune responses during UTI (46–49). Consistent with these roles, female mice mount stronger IL-17-mediated responses and more effectively clear UPEC, whereas males are more prone to chronic infection (23, 30). These observations, together with our findings of sex-specific transcriptional responses with obesity, suggest that hormonal context shapes how the bladder urothelium responds to metabolic stress. Collectively, these data support a broader model in which metabolic and hormonal signaling coordinate urothelial differentiation and host defense, and in which sex-based biological differences shape the bladder’s response to both infection and obesity.

Consistent with this model of metabolic and hormonal regulation of bladder defense, our pathway analysis revealed shared up-regulation of focal adhesion, ECM-receptor interaction, and cytoskeletal signaling in both sexes. FAK, a central regulator of these pathways, was hyperactivated in urothelium of HFD-fed mice, facilitating UPEC invasion. Prior work using bladder cancer models has shown that UPEC exploits FAK signaling to trigger actin rearrangement and host cell entry, particularly through type 1 pili and the FimH adhesin, which engages with β1-integrins to activate FAK (50, 51). Our findings extend this mechanism to primary urothelial cells and provide the first direct evidence that FAK overexpression promotes UPEC infection. In contrast, inhibition of FAK, whether through genetic approaches or natural compounds, reduced UPEC internalization (50–54). These results position FAK as a key node integrating metabolic stress, epithelial remodeling, and microbial invasion.

An important and somewhat unexpected finding was the striking transcriptional divergence between male and female mice in response to obesity. Despite shared pathway enrichment, the differentially expressed genes were largely non-overlapping between male and female mice. This degree of divergence was surprising, as both sexes displayed increases in UTI susceptibility and FAK activation, suggesting a conserved phenotype despite distinct transcriptional programs. These differences likely reflect broader sex-based variation in metabolic and immune responses to obesity (27, 55–58). While the mechanistic links between obesity, immunity, and tissue-specific remodeling are still emerging, our findings suggest that obesity engages distince transcription responses in males and females that ultimately increase UPEC vulnerability. Moreover, they highlight the importance of including both sexes in studies of UTI defense and support the development of sex-specific strategies for managing infection risk in the context of obesity.

This study has limitations. While the HFD model is well-suited for mechanistic investigation and mirrors key features of human obesity, it does not fully capture the complexity of human disease – including genetic, behavioral, and dietary variation (25, 26). Other obesogenic factors such as high sugar, salt, or saturated fat intake may exert distinct effects on immune and epithelial responses. Microbiota composition and environmental exposures could also influence host-pathogen interactions and UTI susceptibility (2, 59–61). Although we demonstrate a functional role for FAK *in vitro*, we have not yet tested whether pharmacologic inhibition of FAK *in vivo* can reduce bacterial burden or restore epithelial integrity in obesity. Finally, while we observed transcriptional changes in urothelial barrier genes, the precise relationship between FAK activation, urothelial differentiation, and epithelial remodeling remains to be defined. Future studies should investigate how metabolic and hormonal cues converge on FAK signaling to regulate bladder defense across diverse physiological and clinical contexts.

Despite these limitations, our study has strengths. We evaluated both male and female mice, revealing sex-specific transcriptional responses that are often overlooked in preclinical models of UTI and obesity (27). While few studies have examined immune function differences between males and females in the context of obesity, our findings highlight how obesity shapes urothelial and immune responses in a sex-dependent manner. By integrating *in vivo* UTI models, transcriptomics, and *in vitro* mechanistic assays, we provide a comprehensive view of how obesity disrupts bladder defense. This multifaceted approach provides a mechanistic framework linking metabolic stress to impaired bladder defense and supports the development for sex-specific therapeutic strategies to be implemented into clinical practice.

In conclusion, this study identifies focal adhesion signaling as a key pathway by which obesity increases UTI susceptibility. FAK integrates metabolic and mechanical cues to promote epithelial remodeling and UPEC invasion, making it a promising therapeutic target. Future work can explore whether modulating FAK activity can reduce infection burden in obesity. Understanding how metabolic and hormonal cues converge on epithelial signaling networks will be essential for developing precision strategies to protect at-risk patients.

## MATERIALS AND METHODS

### Materials

Details of the reagents used can be found in **Supplemental Table 2**.

### Mice

All mouse experiments were conducted in compliance with the rules and regulations of the Institutional Animal Care and Use Committee. Mice were housed under controlled temperature and humidity conditions, maintained on a 12-hour light-dark cycle, and were provided *ad libitum* access to rodent chow and water. Following weaning, mice were fed Teklad 6% fat standard chow (Inotiv, West Lafayette, IN, USA) until five to six weeks of age. Male and female C57BL/6 mice were then assigned to either a HFD consisting of 60% kilocalories from fat (Research Diets, Inc., New Brunswick, NJ, USA) or SC providing 10% kilocalories from fat (Research Diets). Mice were maintained on a HFD or SC for 4, 8, or 12 weeks.

### Mouse Glucose Measurements and Cytokine Profiles

Blood glucose concentrations were obtained using AlphaTrak glucose monitoring system (Abbott Point of Care, Princeton, NJ, USA). Urinary glucose measurements were assessed by dipstick urinalysis (Chemstrip 2 GP, Roche, Basel, Switzerland). Glucose tolerance testing was performed following a six hour fast using established protocols (62). Blood glucose concentrations were measured using the AlphaTrak glucose monitoring system (Abbott Point of Care, Princeton, NJ, USA) at 15, 30, 60, and 90 minutes post injection of intraperitoneal D-glucose injection. The concentrations of 6 mouse serum cytokines, including IFN-γ, IL-1β, IL-6, IL-10, IL-17A, and TNFα, were measured using a multiplex bead array platform (Bio-Plex, Bio-Rad, Hercules, CA).

### Experimental UTI

Female mice were infected by transurethral catheterization with 10^7^ colony-forming units (CFU) of UPEC (strain UTI89) following established protocols (43, 63). UTI89 is type 1-piliated UPEC strain isolated from a patient with cystitis (64). Male mice were infected by direct bladder inoculation with 10^7^ CFU of UTI89 (46). At the indicated time points, mice were anesthetized and sacrificed via cervical dislocation. Urine was collected, organs were aseptically harvested, and UPEC colonies were enumerated as previously described (43, 63).

### Flow Cytometry

Bladders were harvested 24-hours after experimental UTI with GFP-expressing UTI89. Tissues were dissociated, single live cell suspensions were prepared, and stained for cell surface receptors with fluorescent monoclonal antibody combinations as previously published (**Supplemental Table 2**) (44, 65). UPEC+ cells were gated to enumerate phagocytosis by specific immune cell populations. Intracellular killing was evaluated by mean fluorescence intensity of intracellular GFP-UPEC. Stained cells were collected on a LSR II cytofluorometer (BD Biosciences, Franklin Lakes, NJ, USA) and data analyzed using the FlowJo software as previously published (Treestar, Ashland, TN, USA) (44).

### PMN Isolation and *Ex vivo* UPEC Killing

Neutrophils were isolated from mouse femurs and tibias using the Mojosort Mouse Neutrophil Isolation Kit (BioLegend) following established protocols and were resuspended in RPMI medium with 10% fetal bovine serum (2 x 10^5^ cells/200μl) and placed in 96 well plates at 37°C (65, 66). PMNs were infected with UPEC at an MOI 100 or 1,000 at 37°C. After a 45 minute incubation, supernatants were collected after 45 minutes to assess the extracellular bacterial load. Samples were serially diluted and plated on LB Agar plates. UPEC CFUs were quantified the following day.

### Urothelium Enrichment and RNA Sequencing

Bladders were aseptically harvested from mice maintained on either SC or HFD. Bladders were everted under a dissecting microscope using sterile forceps, then incubated at 37°C for 45 minutes in 2.5 mg/mL Dispase II (Thermo Fisher Scientific, Waltham, MA, USA) prepared in 50 mM HEPES/KOH pH 7.4 with 150 mM NaCl. The urothelium was then mechanically stripped from the detrusor muscle using sterile forceps (44). Isolated cells were washed with cold sterile phosphate buffered saline (PBS) and pelleted for RNA extraction. RNA was extracted using NucleoSpin RNA Plus XS Kit (Takara Bio, San Jose, CA, USA). RNA integrity was assessed using the Agilent Bioanalyzer 2100 (Agilent, Carlsbad, CA, USA). Samples with RNA integrity scores > 7 were submitted for sequencing.

For limiting-cell RNA-seq, RNA was pre-amplified using Clontech SMARTer v4 kit (Takara Bio) and cDNA libraries were generated using Nextera XT DNA Library Prep Kit (Illumina, San Diego, CA, USA). Samples were sequenced to 15 to 20 million paired-end (2 x 150 base pair) clusters on the Illumina HiSeq 4000 platform. Data processing and analysis were performed as previously described (67, 68). The reported sequences and metadata reported are deposited in the Gene Expression Omnibus (GEO) database (Accession No. GSE294660). Differentially expressed gene lists are provided as **Supplemental Dataset 1** (12 weeks on diet) and **Supplemental Dataset 2** (8 weeks on diet). RNA-seq analysis is outlined in the **Supplemental Methods**.

### Cell Culture and Transfection

Primary human bladder urothelial cells (HBLAK), obtained from an 80-year old male, were purchased from CELLnTEC Advanced Cell Systems and grown in CELLnTEC CnT-Prime Epithelial Proliferation Medium (CELLnTEC, Bern, Switzerland) (69, 70). Cells were cultured at 37°C and 5% CO_2_. Transfection experiments were initiated when cells approached 95% confluency. **siRNA Transfection**: Pooled siRNA libraries targeting human *PTK2* and a non-targeting pool (negative control) were purchased (**Supplemental Table 2**). Cells were transfected with a mixture of siRNA, DharmaFECT transfection reagent (Horizon Discovery, Waterbeach, UK), and culture media per the manufacturer’s recommendations. Seventy-two hours after transfection, cells lysates were collected for Western blot or UPEC attachment and invasion assays were performed. **Retrovirus Transduction**: FAK was overexpressed by retroviral transduction of cells with pWZL-Neo-Myr_Flag-DEST (empty control) or pWZL-Neo-Myr-Flag-PTK2 (Addgene, Watertown, MA, USA) (71). Retrovirus was produced using Phoenix-AMPHO cells (ATCC, Manassas, VA, USA). **Plasmid Transfection**: AKT was overexpressed by transfecting cells with 500 ng pCMV5-hemagglutinin (HA)-PKBα (WT) or pCMV5-HA-membrane targeted-PKBα (M/P) (purchased from D. Alessi, University of Dundee, Dundee, United Kingdom) using Lipofectamine 2000 according to the manufacturer’s instructions (Thermo Fisher Scientific, Waltham, MA, USA) (72). For all experiments, culture media was changed 16 hours after transfection or transduction. Experiments were performed 72 hours post transfection or transduction.

### *In vitro* UPEC Infection Assays

UPEC invasion assays were performed as previously described using HBLAK cells (44, 73). Seventy-two hours following transfection or transduction, cells were infected with 10 multiplicity of infection (MOI) UTI89. After 2 hours, one set of wells was lysed in 0.1% Triton X-100 to enumerate total UPEC per well (extracellular and intracellular). To define UPEC invasion, a second set of wells were incubated in culture medium supplemented with gentamicin to kill extracellular bacteria. After 3 hours, cells were washed and lysed to evaluate bacterial invasion. Lysates were plated and bacterial CFUs were enumerated after overnight growth on LB agar. The percentage of intracellular bacteria was calculated as the number of UPEC recovered divided by the total number of UPEC (44, 73).

### Western Blotting and Quantification

Mouse urothelium, total bladder, and HBLAK tissue culture lysates were prepared by homogenizing tissue or cells in RIPA buffer (25 mM Tris-HCl, pH 7.6; 150 mM NaCl, 1% NP-40, 1% sodium deoxycholate, 1% sodium dodecyl sulfate) supplemented with protease and phosphatase inhibitor (Thermo Fisher Scientific, Waltham, MA, USA). Western blot was performed as previously described (73). Antibodies directed against the following targets were used P-FAK (Y397), P-AKT (S473), GAPDH (Cell Signaling, Danvers, MA, USA); M2-FLAG tag, HA tag (Sigma Aldrich, Burlingame, MA, USA). Quantitation of Western blot intensity was performed using ImageLab software (Bio-Rad, Hercules, CA, USA). Relative expression was determined by dividing the background-corrected intensity of P-FAK or P-AKT band by the background-corrected intensity of corresponding GAPDH band.

### Immunostaining and Microscopy

Immunofluorescent staining was performed using published methods (44, 73). Bladders were labeled with a rabbit anti-FAK antibody (Abcam, Waltham, MA, USA), polyclonal goat anti-Uroplakin 3 (Upk3) antibody (Santa Cruz Biotechnology, Dallas, TX, USA). Alexa Fluor® 488 AffiniPure donkey anti-goat IgG and Cy™3 AffiniPure F(ab’)₂ fragment donkey anti-rabbit IgG served as secondary antibodies (Jackson ImmunoResearch Laboratories, West Grove, PA, USA). Nuclei were labeled using Hoechst (Thermo Fisher Scientific, Waltham, MA, USA). Microscopy images were captured using a Nikon Ti2-E microscope and DS-RI2 camera (Nikon Instruments Inc., Melville, NY, USA).

### STATISTICAL ANALYSIS

Continuous differences between groups were evaluated for a normal distribution with the D’Agostino-Pearson Omnibus or Shapiro-Wilk normality test, with normality defined as a *P-*value > 0.05. Comparisons on normally distributed data were performed using a Student’s *t*-test or ANOVA; otherwise, the nonparametric Mann-Whitney *U* or Kruskal-Wallis tests were used. Data from *in vitro* experiments were normally distributed and are presented as means +/- SEM. Differences between groups with a *P-*value < 0.05 were considered statistically significant.

## Acknowledgments and Funding Sources

We thank Birong (Rollin) Li in the Kidney and Urinary Tract Center at the Abigail Wexner Research Institute at Nationwide Children’s Hospital for assistance with the mouse infections. This work is supported by the National Institutes of Health (NIDDK) K01 DK128379 (J.R-R.), R01 DK114035, and R01 DK128088 (J.D.S.). Limiting-cell RNA-seq was performed at The Ohio State University Comprehensive Cancer Center’s Genomics Shared Resource which is supported by a Comprehensive Cancer Support Grant P30CA016058.

## Author Contributions

L.S. and J.D.S. supervised the project and contributed to the design and interpretation of all experiments. K.S. managed the mouse colony and completed the in vivo mouse experiments. K.S., A.S., P.J., and J.R-R. performed mouse biometric testing, flow cytometry, or Western blotting. X.W. and L.S. analyzed the limiting cell RNA-seq. L.S., S.L.P., and A.R.J. completed immunostaining and microscopy. L.S., I.C-O., and Y.S-Z. performed the ex vivo and in vitro experiments. L.S. and J.D.S. wrote the manuscript with input from all authors.

## Supporting Information for

### Supplemental Methods

#### Mouse Limiting Cell RNA-seq Analysis

##### RNA-seq data preprocessing, alignment, and quantification

Individual FASTQ files were trimmed for adapter sequences and filtered for a minimum Phred quality score of Q20 using Adapter Removal v2.2.0 – indicating that the base call accuracy is 99% or greater (1). Preliminary alignment was performed using HISAT2 to a composite reference of ribosomal RNA, mitochondrial DNA, and PhiX bacteriophage sequences obtained from National Center for Biotechnology Information Reference Sequence Database (RefSeq) (2). Reads aligning to these references were excluded in subsequent analyses. Primary alignment was performed against the mouse genome reference GRCm38p4 using HISAT2 alignment program. Gene expression values for genes described by the GENCODE Gene Transfer Format release M14 were quantified using the featureCounts tool of the Subread package in unstranded mode (3–6).

##### Sequencing and alignment quality control

Quality control was performed using a modification of the QuaCRS custom workflow (7). In brief, aligned read quality was verified using RNA-SeQC and RSeQC (8, 9). The exonic rate (the percentage of reads aligning to exons), intronic rate (the percentage of reads aligning to introns), and duplication rate (the percentage of reads that were identified as PCR duplicates) were evaluated.

##### Coverage profiling

Coverage depths across the aligned reference were calculated with a per-base resolution using the ‘genomecov’ utility of Bedtools in BedGraph format (10). These files were used as input into CLEAR: a gene-by-gene analysis workflow to identify genes in ultra-low input RNA-seq data for downstream analyses. Determination of analysis-ready CLEAR transcripts was performed as previously described (11).

##### RNA-Seq data analysis

Principal component analysis (PCA) visualized differences between samples. All PCA plots were generated from counts tables that were size-normalized and r-log transformed after CLEAR selection using methods included with DESeq2. Comparisons were processed using plotPCA and visualized using ggplot (12–14). Differentially expressed genes were called using DESeq2 from CLEAR counts tables generated with featureCounts as described above (12). In all figures, a false discovery rate *q*-value of < 0.05 was used as the inclusion criterion for differentially expressed genes (DEGs). DEGs were visualized by volcano plot generated using ggplot.

##### Venn diagram analysis

Overlapping upregulated or downregulated DEGs from female and male mice were determined using Biovenn.(15) Venn diagrams were made in Microsoft PowerPoint.

##### KEGG pathway analysis

PathfindR was used to identify KEGG pathways enriched from DEGs with adjusted *P-*value < 0.05.(16) Upregulated genes (log2FoldChange > 0) and downregulated (log2FoldChange < 0) DEGs were used as input, separately. Pathway enrichment was reported as a bubble plot made using ggplot; size of bubbles represented the number of DEGs in each term and color represented *P*-value of enrichment. Differential expression of individual genes within KEGG terms were visualized by bubble plot generated using ggplot.

## Supplemental Figures and Figure Legends

**Supplemental Figure 1:**
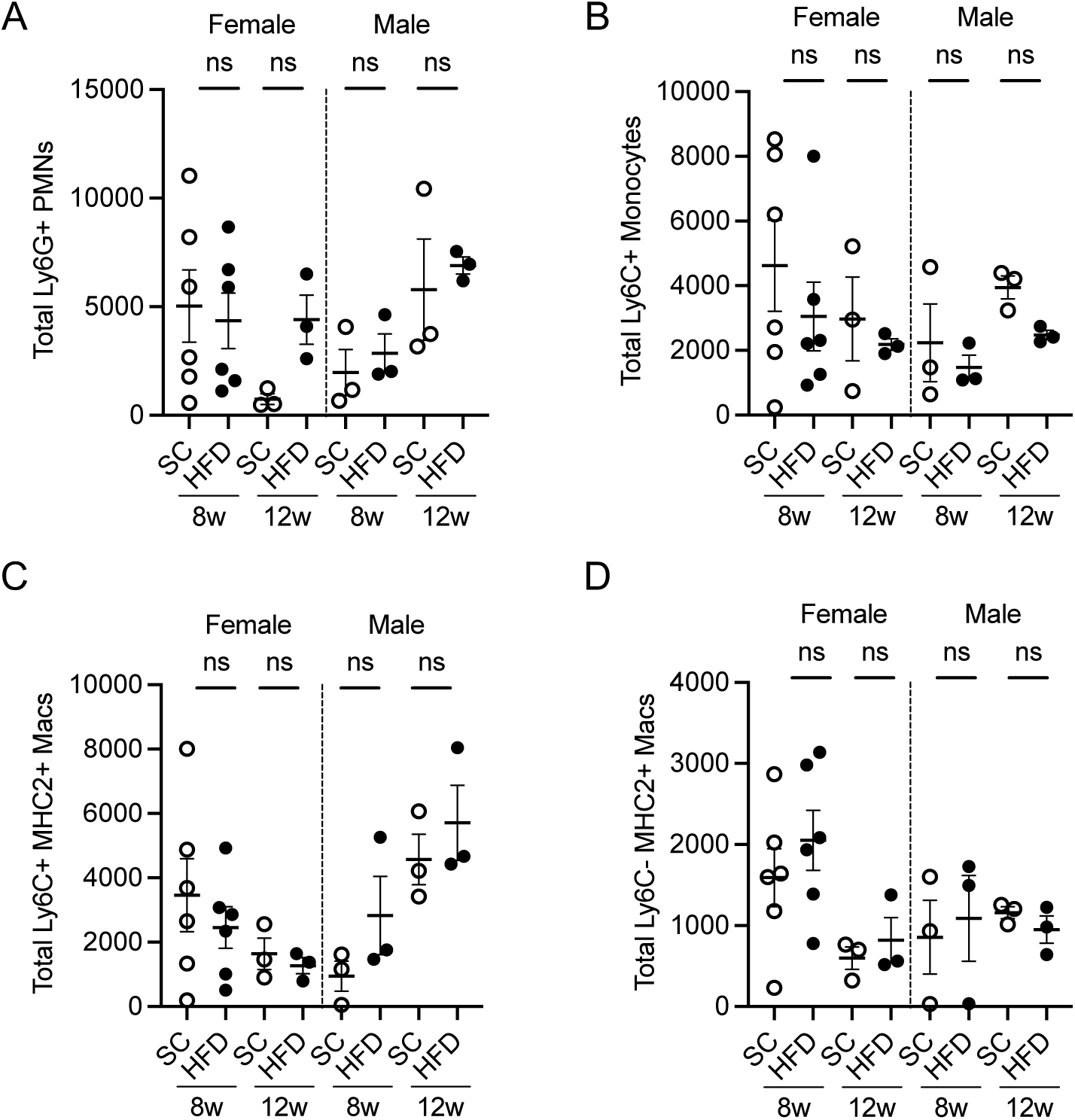
Characterization of immune cell recruitment in UPEC-infected bladders. Female and male mice maintained on a HFD or SC were infected with UPEC and mouse bladders were collected for flow cytometry twenty-four hours after infection. (**A-D**) Plots of the absolute numbers of neutrophils (A), inflammatory monocytes (B), monocyte-derived macrophages (C), and resident macrophages (D) in infected bladders. Graphs show the mean and SEM. Each point denotes immune cell numbers in a unique mouse (*n*=3-6). Pairwise comparisons were performed between HFD and SC groups as indicated in graphs (Mann Whitney *U* test). **These data supplement Figures 1 and 2.**

**Supplemental Figure 2:**
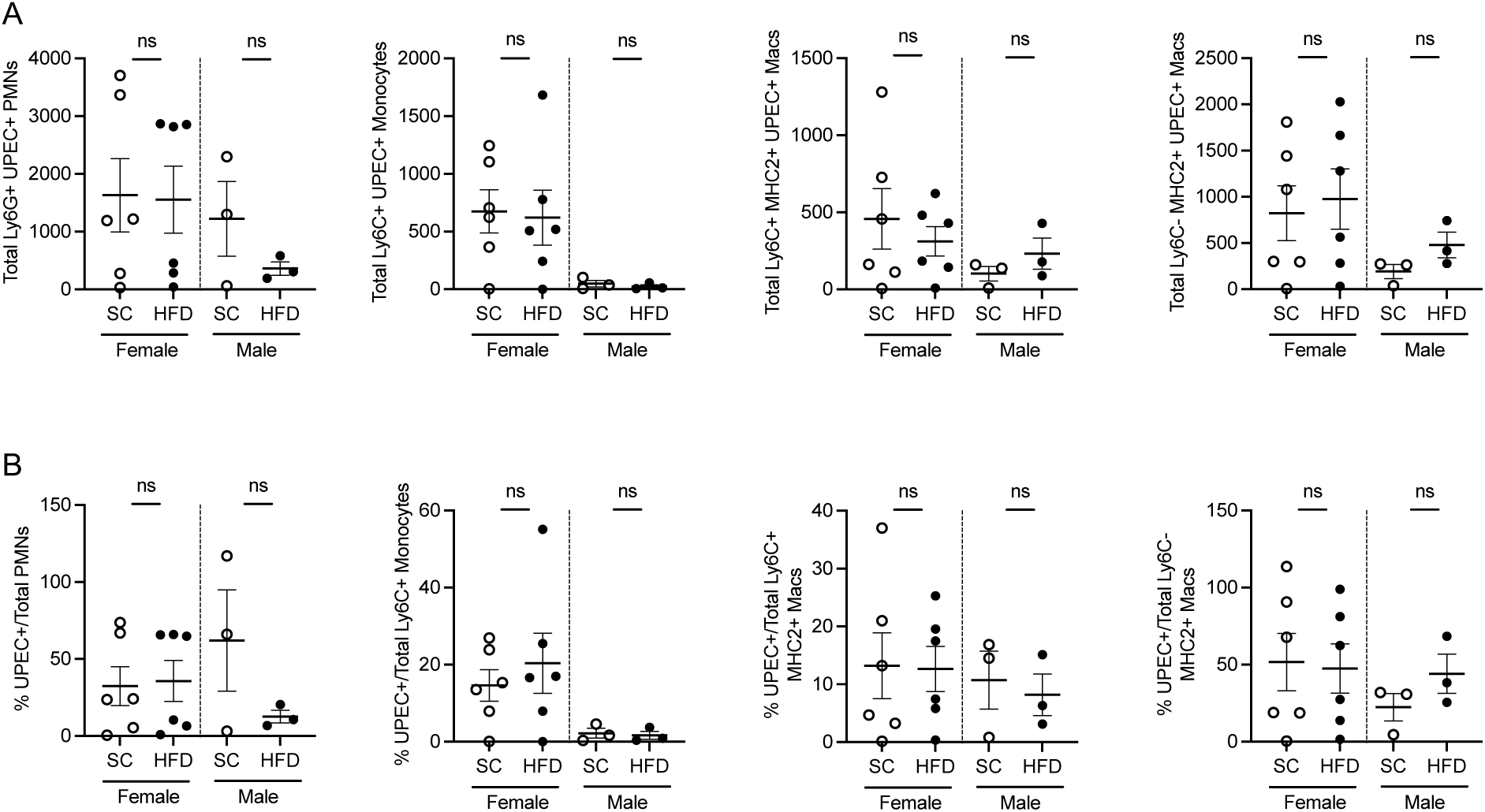
Diet-induced obesity does not impact UPEC phagocytosis. Female and male mice were maintained on a high-fat diet (HFD) or standard chow (SC) for 8 weeks, then infected with GFP-expressing UPEC. Bladders were collected twenty-four hours after infection for flow cytometry. (**A**) Plots of absolute numbers UPEC-positive neutrophils, monocytes, monocyte-derived macrophages, and resident macrophages, indicative of UPEC phagocytosis by these myeloid populations (**B**) Percent of UPEC+ cells within each immune cell population, reflecting the proportion of phagocytic activity. Graphs show the mean and SEM. Each point denotes immune cell numbers in a unique mouse (*n*=3-6). Pairwise comparisons were performed between HFD and SC groups as indicated in graphs (Mann Whitney *U* test). **These data supplement Figures 1 and 2.**

**Supplemental Figure 3:**
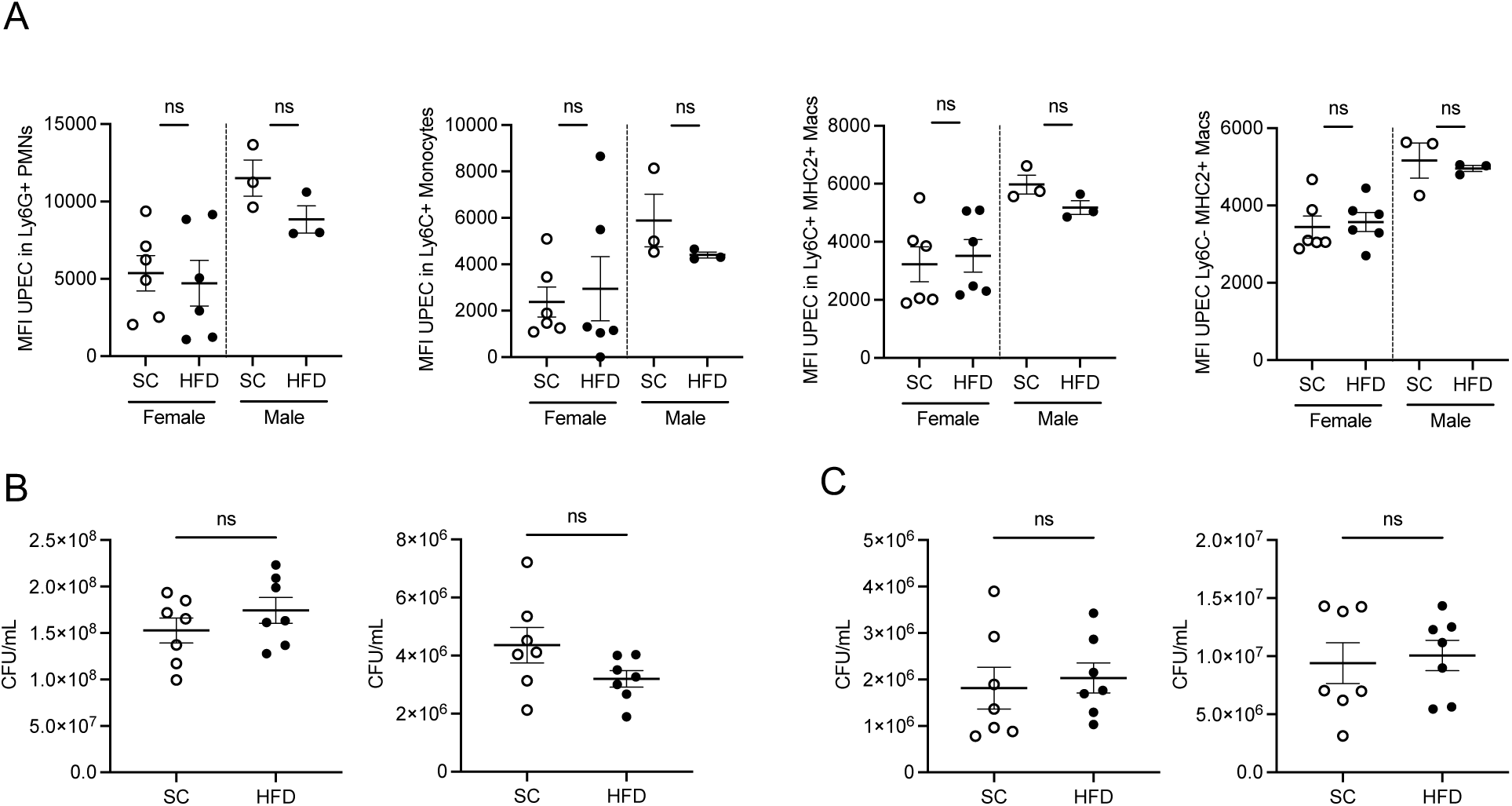
Diet induced obesity does not impact intracellular or extracellular UPEC killing. (**A**) Female and male mice maintained on a high-fat diet (HFD) or standard chow (SC) for 8 weeks, then infected with GFP-expressing UPEC. Bladders were collected twenty-four hours after infection for flow cytometry. Plots illustrate mean fluorescence intensity (MFI) of GFP+ signal in neutrophils, monocytes, monocyte-derived macrophages, and tissue-resident macrophages. MFI serves as a surrogate to measure intracellular UPEC burden, with higher MFI reflecting increased UPEC survival or impaired killing. (**B**-**C**) Neutrophils were enriched from bone marrow of uninfected female mice maintained on a HFD or SC diet for 8 weeks, then challenged *ex vivo* with UPEC strain UTI89 (B) or CFT073 (C) at 100 MOI (left) or 1000 MOI (right). Following incubation, extracellular bacterial killing was enumerated and graphed as CFU/mL. Graphs show the mean and SEM. Each point denotes a direct measurement from an individual mouse (A) or neutrophils recovered from a unique mouse (B-C; *n*=3-7). Pairwise comparisons were performed between HFD and SC groups using the Mann Whitney *U* test. **These data supplement Figures 1 and 2.**

**Supplemental Figure 4:**
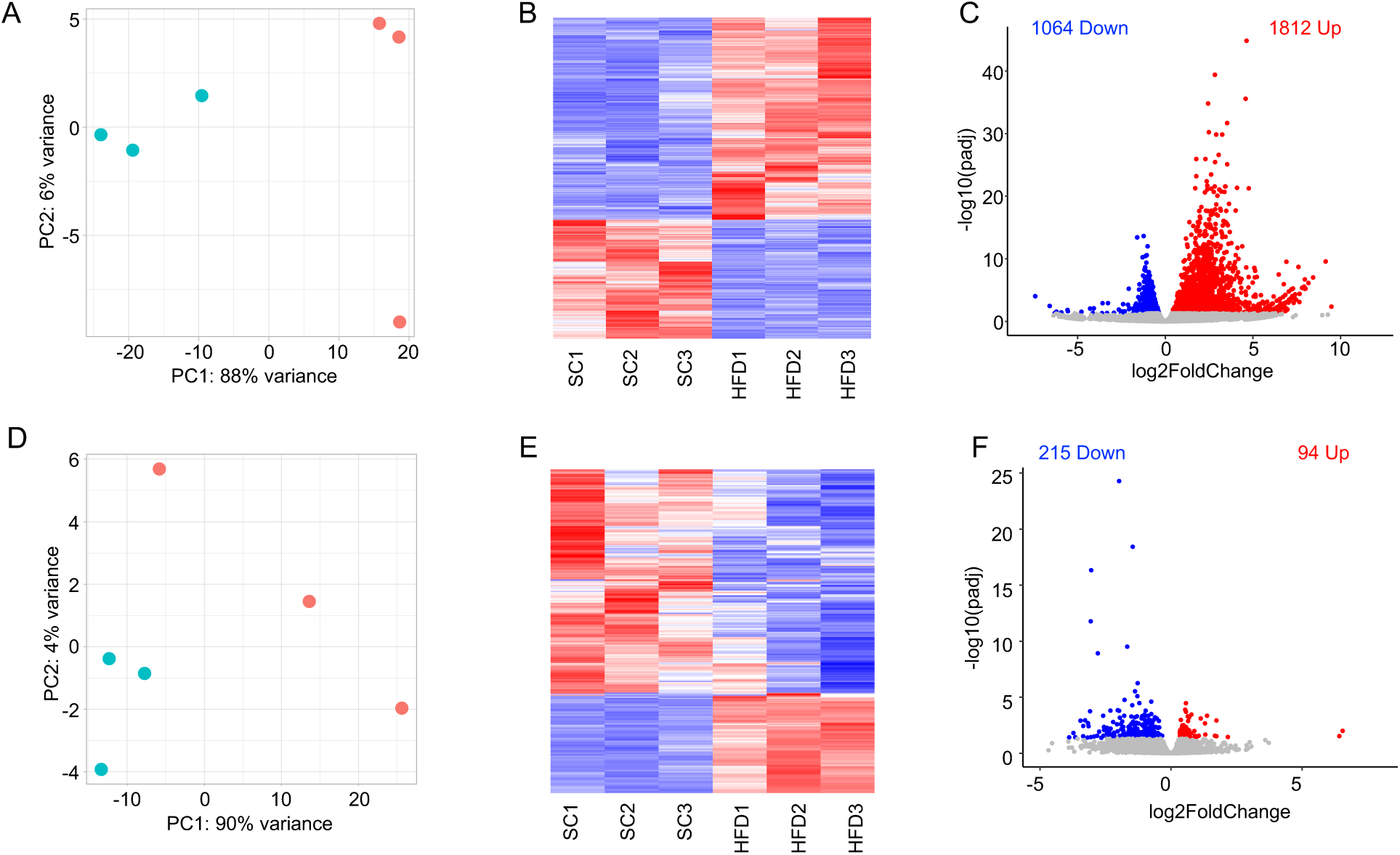
Eight weeks of high-fat diet alters gene expression in the female and male urothelium. (**A/D**) Principal component analysis (PCA) plots from RNA-seq data of female (A) and male (D) urothelium after 8 weeks on standard chow (SC) or high-fat diet (HFD). Each symbol represents a urothelial sample from an individual mouse fed SC (green) or HFD (red). (**B**/**E**) Heatmaps showing differential gene expression between HFD and SC fed female (B) and male (E) urothelium, with each column representing an individual mouse. (**C**/**F**) Volcano plots illustrating differentially expressed genes expression between HFD and SC fed female (C) and male (F) urothelium. Red marks up-regulated genes with log2 fold change > 0 and adjusted *P*-value < 0.05. Blue marks down-regulated genes with log2 fold change < 0 and adjusted *P*-value < 0.05. Adjusted *P*-values were calculated using the Benjamini-Hochberg procedure. **These data supplement Figure 3.**

**Supplemental Figure 5:**
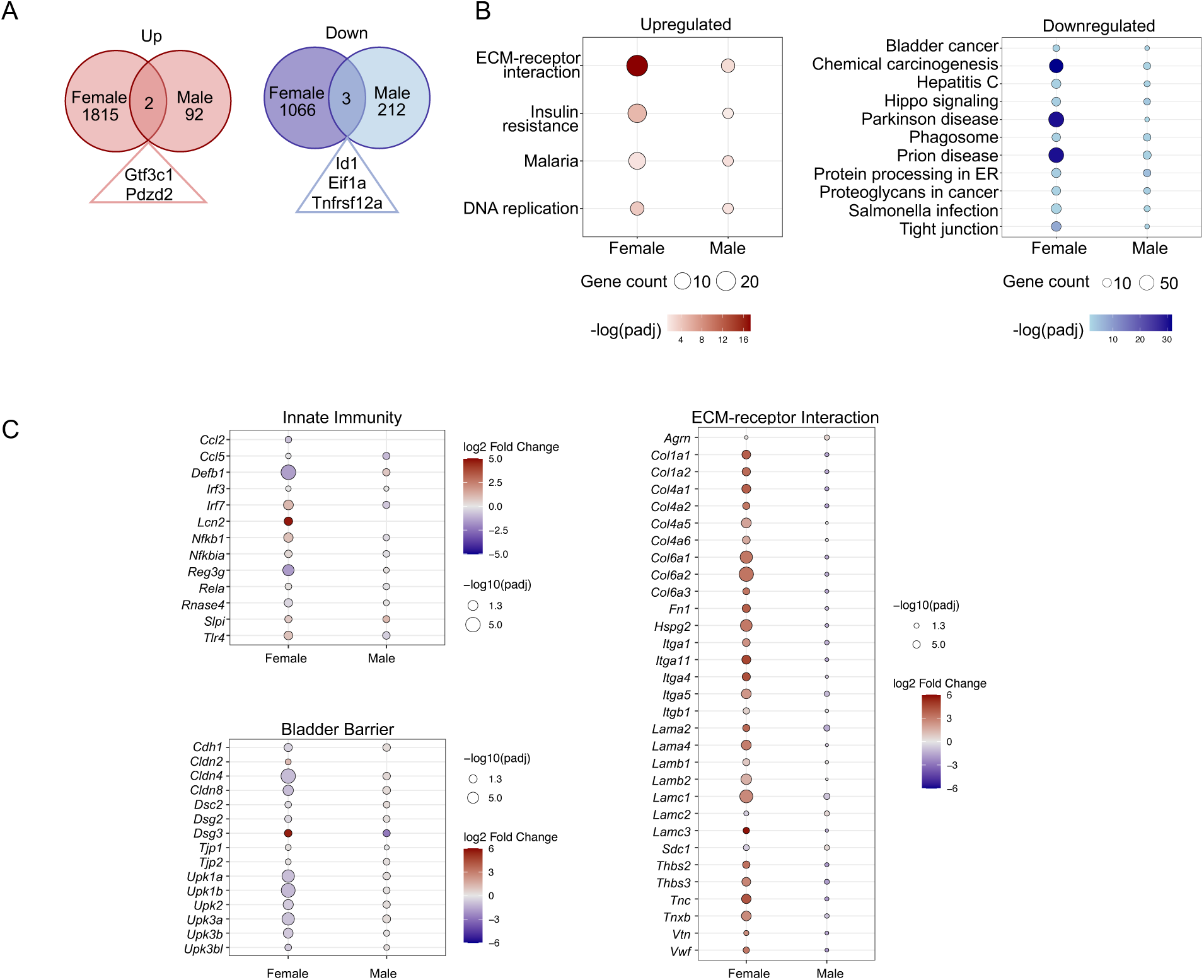
Eight weeks of high-fat diet consumption leads to increased cell-ECM signaling in the urothelium. (**A**) Venn diagrams show minimal overlap between significantly up-regulated (left) and down-regulated (right) genes in female and male urothelium after 8 weeks on standard chow (SC) or high-fat diet (HFD). (**B**) KEGG pathway enrichment derived from up-regulated (left) and down-regulated (right) genes in HFD-fed female and male mice. Bubble size reflects the number of enriched genes per pathway and color indicates significance. (**C**) Expression of curated gene sets related to innate immunity, bladder barrier integrity, and ECM-receptor interaction. Bubble color represents log2 fold change (HFD vs. SC) and size indicates significance. (B-C) Significance was assessed via hypergeometric testing on active subnetworks identified from the protein-protein interaction network using PathfindR, with *P*-values adjusted using the Benjamini-Hochberg method. **These data supplement Figure 4.**

**Supplemental Figure 6:**
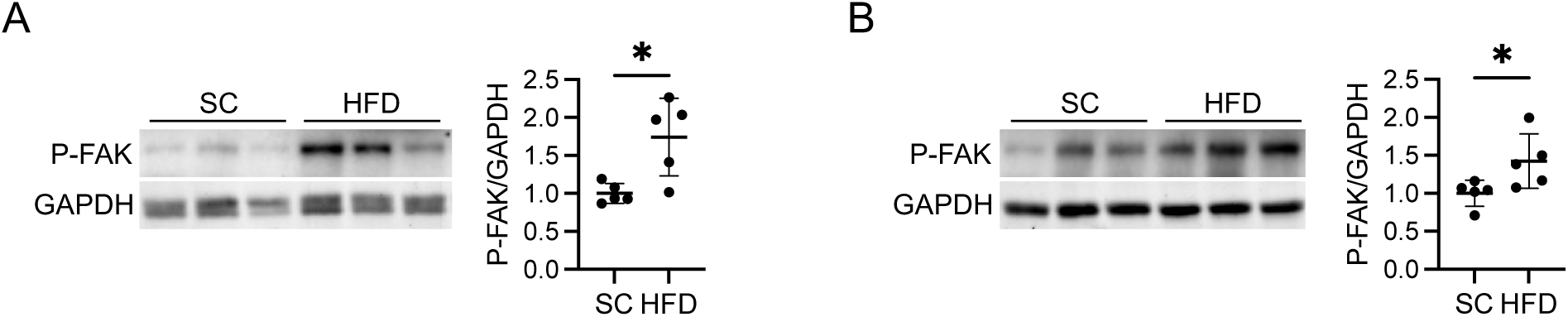
Focal adhesion kinase signaling is elevated in total bladders during obesity. (**A/B**) Representative Western blots (left) and densitometry (right) of P-FAK (Y397) and GAPDH in total bladder lysates from female (A) and male (B) mice maintained on standard chow (SC) or high-fat diet (HFD) for 8 weeks. Quantification of Western blots was performed on bladders from 5 mice per group (*n* = 5). Graphs show the mean and standard deviation. Each point represents an individual mouse. Asterisks indicate significant P-values for the indicated pairwise comparisons (Mann-Whitney *U* test). *P < 0.05. **These data supplement Figure 5.**

**Supplemental Figure 7:**
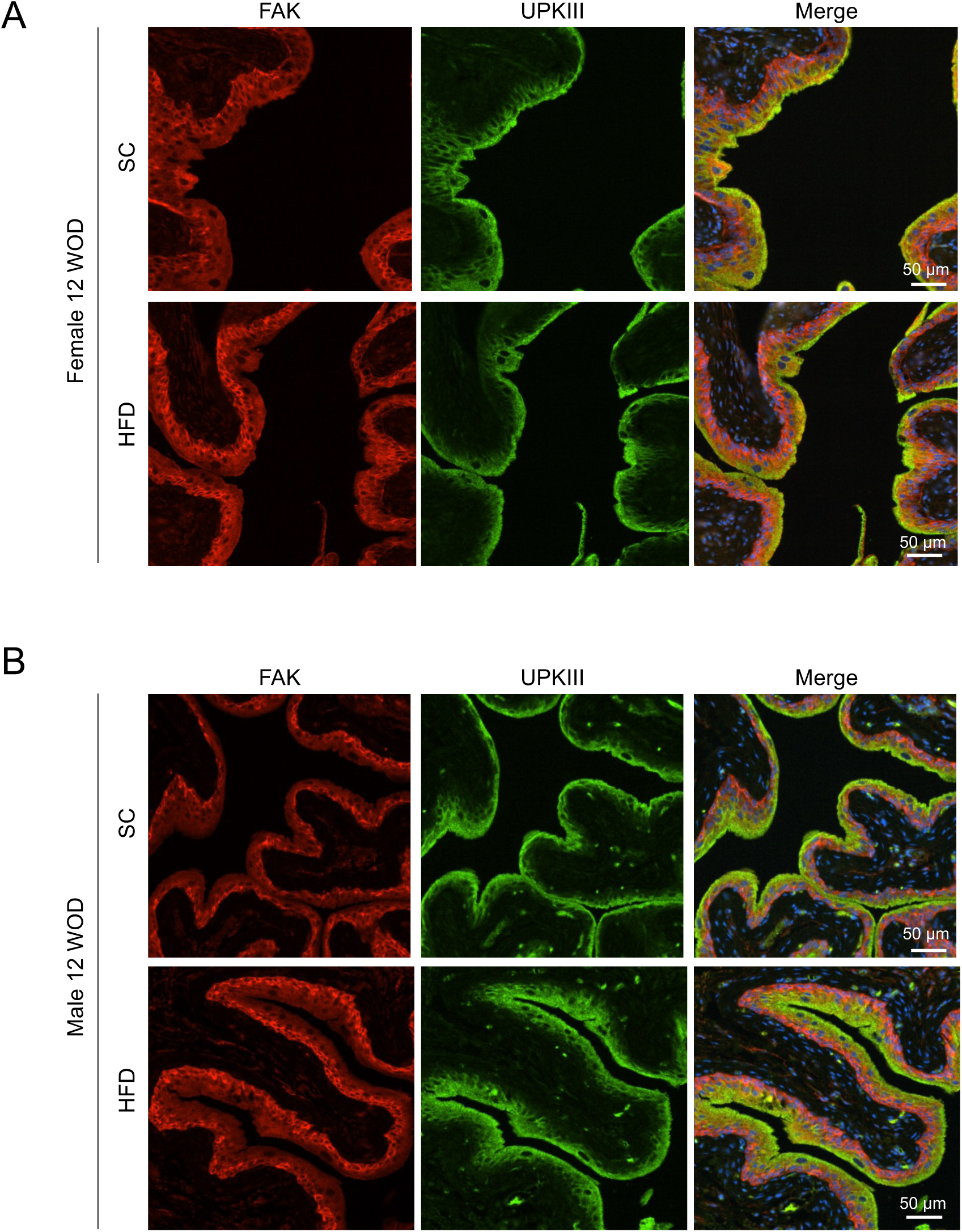
Obesity increases urothelial FAK expression. Representative immunofluorescent bladder images from non-infected (A) female and (B) male mice after 12 weeks on standard chow (SC) or high-fat diet (HFD) labeled with antibodies to total FAK (red), uroplakin III (green) and nuclei (blue, Hoechst). Magnification = 20X. Staining was completed on 4 bladders per cohort. **These data supplement Figure 5.**

## Supplemental Tables

**Supplemental Table 1:** Cytokine profiles in diet induced obesity male and female mice. Serum cytokine profiles in female and male mice fed standard chow (SC) or a high-fat diet (HFD) for 12 weeks (*n* = 7 mice/cohort). *P*-values evaluate differences between sex-matched cohorts (Mann-Whitney *U* test). IL-1β was below the limits of detection (LOD).

|  | Female |  |  | Male |  |  |
| --- | --- | --- | --- | --- | --- | --- |
|  | SC | HFD |  | SC | HFD |  |
| Cytokine (pg/mL) | Mean ± SEM | Mean ± SEM | *P*-value | Mean ± SEM | Mean ± SEM | *P*-value |
| IFN-γ | 22.05 ± 1.21 | 25.63 ± 1.28 | 0.073 | 30.62 ± 2.68 | 27.84 ± 4.58 | 0.277 |
| IL-1β | < LOD | < LOD | NA | < LOD | < LOD | NA |
| IL-6 | 14.24 ± 0.35 | 15.89 ± 0.91 | 0.277 | 18.16 ± 1.17 | 19.42 ± 3.17 | 0.594 |
| IL-10 | 69.58 ± 1.78 | 81.6 ± 3.04 | **0.004** | 81.6 ± 2.25 | 79.11 ± 3.94 | 0.435 |
| IL-17A | 40.5 ± 8.57 | 70.77 ± 12.87 | 0.073 | 102.3 ± 18.38 | 58.49 ± 12.51 | 0.128 |
| TNFα | 78.77 ± 10.07 | 96.34 ± 6.66 | 0.104 | 130.3 ± 20.69 | 94.32 ± 11.85 | 0.165 |

**Supplemental Table 2:** Key Reagents. List of mice, antibodies, chemical reagents, recombinant plasmids, and siRNA used with vendor and catalog number.

| Reagent | Vendor | Catalog Number |
| --- | --- | --- |
| Mice |  |  |
| B6.Cg-Gt(ROSA)26Sor<sup>tm14</sup>(CAG-tdTomato)<sup>Hze</sup>/J | The Jackson Laboratory | Strain # 007914 |
| Diet |  |  |
| Teklad Chow | Inotiv | 2920X |
| Standard Chow (SC) | Research Diets, Inc. | D12450Ji |
| High-Fat Diet (HFD) | Research Diets, Inc. | D12492i |
| Cell Culture |  |  |
| Primary human urothelial cells (HBLAK) | CELLnTEC Advanced Cell Systems | HBLAK<br>RRID:CVCL_JQ59 |
| CnT-Prime Epithelial Proliferation Medium | CELLnTEC Advanced Cell Systems | CnT-PR |
| Phoenix-AMPHO cells | American Type Culture Collection (ATCC) | CRL-3213 |
| Bacteria |  |  |
| Uropathogenic *Escherichia coli* UTI89 | Provided by Dr. Scott J. Hultgren | PMID: 2877947 |
| Plasmids |  |  |
| pWZL-Neo-Myr-Flag-DEST (empty) | Addgene | 15300 |
| pWZL-Neo-Myr-Flag-PTK2 | Addgene | 20610 |
| pCMV-HA-PKBα-WT | Provided by Dr. Dario Alessi | PMID: 9395488 |
| pCMV-HA- PKBα-M/P (membrane-targeted) | Provided by Dr. Dario Alessi | PMID: 9395488 |
| Antibodies |  |  |
| GAPDH | Cell Signaling Technology | 2118 |
| P-AKT (Ser473) | Cell Signaling Technology | 4060 |
| P-FAK (Tyr397) | Abcam | ab81298 |
| M2-FLAG tag | Sigma Aldrich | F3165 |
| HA Tag | Sigma Aldrich | 05-904 |
| FAK | Abcam | ab40794 |
| Uroplakin IIIA | Fitzgerald | 10R-U103ax |
| Goat anti-Rabbit (Cy3) | Jackson ImmunoResearch | 111-165-003 |
| Goat anti-Mouse (Alexa Fluor 488) | Jackson ImmunoResearch | 115-545-003 |
| Hoechst | Thermo-Fisher | PI62249 |
| Brilliant Violet 785 anti-mouse CD45 | BioLegend | 103149, 30-F11 |
| Alexa Fluor 700 anti-mouse/human CD11b | BioLegend | 101222, M1/70 |
| Brilliant Violet anti-mouse Ly-6C | BioLegend | 128031, HK1.4 |
| PerCP/Cy5.5 anti-mouse Ly-6G | BioLegend | 127616, 1A8 |
| Chemical Reagents |  |  |
| DharmaFECT transfection reagent | Dharmacon | T-2001-02 |
| Lipofectamine 2000 | Thermo-Fisher | 11668027 |
| Halt Phosphatase and Protease Inhibitor | Thermo-Fisher | PI78445 |
| Commercial Assays |  |  |
| Nucleospin RNA Plus XS Kit | Takara Bio | 740990 |
| Clontech SMARTer v4 Ultra Low Input Kit | Takara Bio | 634888 |
| Nextera XT DNA Library Preparation Kit | Illumina | FC-131-1024 |
| Bio-Plex Pro Mouse Cytokine Th17 Panel A | Bio-Rad | M6000007NY |
| Mojosort Mouse Neutrophil Isolation Kit | BioLegend | 480058 |
| siRNA Pools |  |  |
| SMARTpool: Non-Targeting siRNA Pool | Horizon Discovery | D-001810-10-05 |
| SMARTpool: ON-TARGETplus PTK2 siRNA | Horizon Discovery | L-003164-00-0005 |

**Dataset S1 (separate file): Differentially expressed genes in urothelium after 12 weeks of HFD vs SC diet feeding.** These data supplement Figures 3 and 4.

**Dataset S2 (separate file): Differentially expressed genes in urothelium after 8 weeks of HFD vs SC diet feeding.** These data supplement Supplemental Figures 4 and 5.

## Notes

**Competing Interest Statement:** The authors declared that no conflict of interest exists.

### Competing Interest Statement

The authors have declared no competing interest.

### Summary of Updates

Added significance summary and minor edits for formatting. No new data.

